# Similarity and dissimilarity in alteration of gene expression profile associated with inhalational anesthesia between sevoflurane and desflurane

**DOI:** 10.1101/2023.05.21.541665

**Authors:** Takehiro Nogi, Kousuke Uranishi, Ayumu Suzuki, Masataka Hirasaki, Tina Nakamura, Tomiei Kazama, Hiroshi Nagasaka, Akihiko Okuda, Tsutomu Mieda

## Abstract

Although sevoflurane is the most commonly used inhalational anesthetic agent, the popularity of desflurane is increasing to a similar level. The main beneficial property of desflurane is the relatively fast emergence of the patient from the anesthetic state after halting its supply compared with anesthesia using other anesthetic agents. However, there has been no comprehensive comparison of the effects of these two anesthetic agents on alterations in liver gene expression profiles in animals, including humans, to assess the levels of hepatotoxicity that is induced at least in some extent by inhalational anesthesia. Thus, we compared alterations in the global gene expression profiles in the livers of rats subjected to inhalational anesthesia by sevoflurane or desflurane by a next-generation sequencing method. Our data revealed that both anesthetic agents significantly activated a similar set of genes including those related to drug metabolism and circadian rhythm. Furthermore, many genes downregulated by sevoflurane were also downregulated by desflurane. However, many of the genes related to the cholesterol biosynthetic process were specifically repressed by sevoflurane, but not by desflurane.

## Introduction

Inhalational anesthesia using halogenated anesthetic agents is the most commonly used method for general anesthesia that induces sedation in patients to ensure an appropriate surgical field^1,2^. However, safety issues were a serious concern for the use of halogenated anesthetics^3,4^. Indeed, the utilization of halothane, a classical anesthetic agent, caused hepatic injuries in many cases^5,6^. Halothane is hardly used clinically because of this risk and sevoflurane is currently the most commonly used inhalational anesthetic agent because of its numerous beneficial characteristics in the clinic^7,8^. For example, compared with halothane, sevoflurane is much more refractory to being dissolved in the blood leading to its elimination from the lung as a vapor^9,10^. Furthermore, sevoflurane, unlike other halogenated anesthetic agents, does not produce trifluoroacetic acid, which can induce severe hepatic injury via activation of a patient’s immune response during its metabolism in the liver^11,12^. Because of these remarkable features, sevoflurane is regarded as an extremely safe anesthetic agent. Despite this, desflurane, another halogenated anesthetic agent, is rapidly gaining popularity and the prevalence of inhalational anesthesia using desflurane is approaching that of sevoflurane. Although desflurane has some disadvantages such as respiratory tract irritation^13-17^, an obvious advantage of its clinical use is its relatively rapid induction of an anesthetic state upon administration and the rapid recovery of patients from that state after the cessation of its supply compared with other anesthetic agents^18-23^. These effects of desflurane are related to its marked low solubility in blood, which is even lower than that of sevoflurane^10,24,25^. An additional notable characteristic of desflurane is its extreme resistance to degradation and biotransformation^10,26-28^, further elevating its safety level for clinical use. Thus, both sevoflurane and desflurane have beneficial features as inhalational anesthetic agents. However, it is not known which anesthetic agent causes relatively stronger influences in global expression profile in liver that may be correlated with levels of hepatotoxicity.

In this study, we addressed this question by comparing alterations in the global gene expression profiles of livers in rats subjected to inhalational anesthesia by desflurane or sevoflurane. Consistent with a previous report, we found that general anesthesia using sevoflurane led to the significant activation of genes related to drug metabolizing enzymes and circadian rhythm^29,30^. Our data revealed that general anesthesia using desflurane also activated sets of genes similar to those activated by sevoflurane. Likewise, similar sets of genes were downregulated by sevoflurane and desflurane, indicating these agents cause comparable stress and damage in the liver when used to induce anesthesia. However, as an exception, our data revealed that genes related to the “cholesterol biosynthetic process” were coordinately repressed by sevoflurane but not desflurane.

## Materials and Methods

### Animal experimentation

Male Wister rats (6 weeks old, 140–160 g body weight) purchased from Japan SLC Inc. (Hamamatsu, Japan) were housed in plastic cages with free access to food and water at 23°C under controlled lighting (12:12-hour light/dark cycle: lights on at 7:00 AM) for at least 1 week to acclimatize. Rats were deprived of food and water for 2 hours prior to experimentation and then subjected to inhalational anesthesia via nose cones using sevoflurane (4.5% gas-air mixture)^29^ or desflurane (6.0% gas-air mixture)^27^ with a 3 L/min flow of 50% oxygen. During anesthesia, rats were allowed to breathe spontaneously and the value of SpO_2_ measured using a pulse oximeter through the lower extremities of rats was strictly controlled to not drop below 98%. In addition, the body temperatures of anesthetized rats were maintained at 36.5–37.5°C. After 6 hours, rats subjected to inhalational anesthesia with sevoflurane (Sevo6) or desflurane (Des6) were sacrificed by decapitation and the left lateral lobe of the liver was quickly isolated from each rat. Then, a portion of each liver was immersed in RNA later after rinsing with phosphate-buffered saline and stored at 4°C. For control rat livers, rats subjected to inhalational anesthesia with sevoflurane (Sevo0) or desflurane (Des0) were immediately sacrificed after loss of consciousness and liver specimens were prepared and stored as above. The protocol of these experiments was approved by the Institutional Review Board on the Ethics of Saitama Medical University (permission numbers 3301, 3547, and 3796).

### RNA preparation and reverse transcription

Total RNAs were prepared from livers of rats subjected to inhalational anesthesia with sevoflurane or desflurane for 6 hours or less than 1 minute using an RNeasy Midi Kit (QIAGEN, Venlo, Netherlands) according to the manufacturer’s instructions. RNAs were then used to obtain cDNAs by reverse transcription as described previously^31^.

### Quantitative PCR

cDNAs obtained by reverse transcription were used for quantitative PCR (qPCR) using the following TaqMan probes.

*Alas1*: Rn00577936_m1; *Tp53i13*: Rn01497061_g1; *Cyp2b1*: Rn01457880_m1; *Cyp7a1*: Rn00564065_m1; *Per3*: Rn00709499_m1; *Dbp*: Rn01498425_m1; *Fdps*: Rn00821389_g1; *Fdft1*: Rn00570323_m1; *Por*: Rn00580820_m1; *Hsd17b2*: Rn00577779_m1; *Zfp354a*: Rn00583768_m1; *Igtp*: Rn01751473_m1; *Irf1*: Rn01456791_m1; *Gapdh*: Rn01775763_g1.

All samples were tested in triplicate and the results were normalized to *Gapdh* expression levels.

### RNA sequencing

Libraries were prepared with total RNAs using TruSeq Stranded Total RNA Library Prep Gold Kit from Illumina (San Diego, California) according to the manufacturer. RNA sequencing was performed on an Illumina platform by paired-end 101 bp reads with 40– 60 M reads for each sample.

### Gene Ontology and Gene Set Enrichment Analyses

Gene Ontology (GO) analysis was performed using DAVID web tools (http://david.abcc.ncifcrf.gov). Gene Set Enrichment Analysis (GSEA)^32^ was conducted according to the method described on the GSEA homepage (http://www.gsea-msigdb.org/gsea/index.jsp) using three different platforms of gene sets, “biological process of Gene Ontology”, “Kyoto Encyclopedia of Genes and Genome”, and “Reactome Pathway Database”.

### Statistical Analysis

All data from qPCR were subjected to the Student’s *t*-test to examine statistical significance.

### Accession Number

RNA sequence data used for this study were deposited under accession number GSE230365.

## RESULTS

### Genome-wide expression analyses of livers from rats subjected to inhalational anesthesia

To compare alterations in global expression profiles of livers caused by inhalational anesthetic agents between sevoflurane and desflurane, we conducted comprehensive gene expression analyses of livers from rats subjected to inhalational anesthesia by next-generation sequencing (Fig. S1). Our data revealed that 434 and 350 genes were transcriptionally activated more than 2-fold by sevoflurane and desflurane, respectively (Table S1). Next, these upregulated genes were assigned to GO classification to correlate gene expression changes with overall molecular functions. These analyses yielded 7 and 11 specific GO terms related to sevoflurane (Fig. 1A) and desflurane (Fig. 1B), respectively, whose *P*-values were less than 10^−4^. Notably, the most and second most significant GO terms, marked in pink and green, respectively, were the same for sevoflurane and desflurane. In addition, two GO terms (marked in yellow or light blue) and one GO term (marked in brown) were common and related (located in the same GO term tree), respectively, indicating that sevoflurane and desflurane promote similar biological effects including the circadian rhythm, which was previously linked to inhalational anesthesia^29,30^. Consistent with this notion, a significant number of genes overlapped between these two gene sets (Fig. 1C).

**Figure 1.**
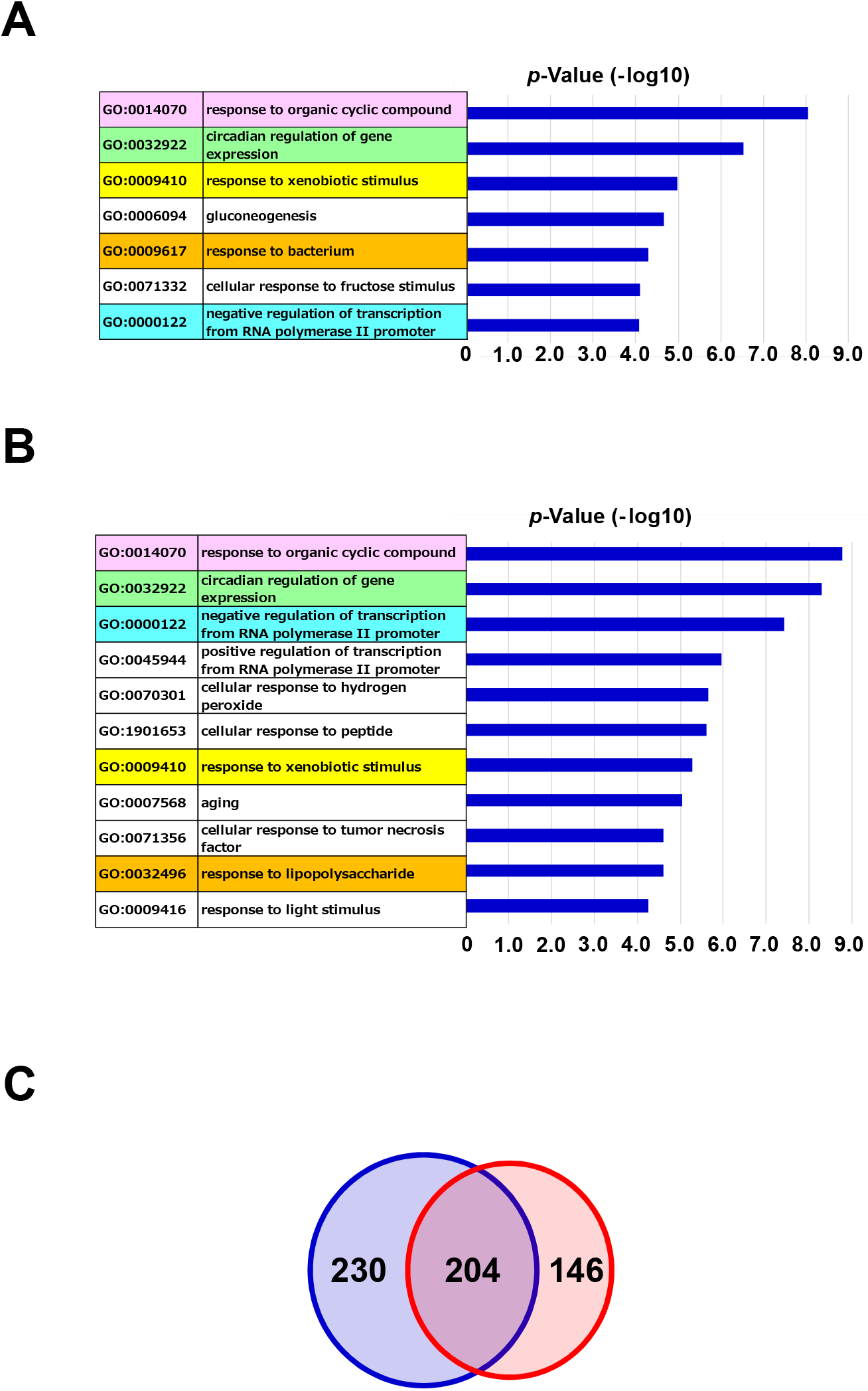
GO analyses of genes upregulated by inhalational anesthesia. Genes whose signal (TPM) values from RNA sequencing increased more than 2-fold after anesthesia induced by sevoflurane (A) or desflurane (B) were individually subjected to GO analyses using DAVID web tools (http://david.abcc.ncifcrf.gov). GO terms with *P*-value less than 10^−4^ were selected and subjected to AmiGo2 analyses (http://amigo.geneontology.org/amigo/landing) to eliminate synonymous terms. GO terms identical or related between sevoflurane and desflurane treatments were marked by distinct colors (pink, light green, yellow, brown, or light blue). (C) Venn diagram showing a comparison of genes upregulated more than 2-fold by sevoflurane or desflurane.

We also conducted GO analyses for genes downregulated by anesthetic agent treatment (Fig. 2A, B, Fig. S1, Table S1). Contrary to the upregulated genes, we found that the analyses of downregulated genes yielded few GO terms (4 and 2 for sevoflurane and desflurane, respectively) with *P*-values less than 10^−4^ and no GO term was shared between the sevoflurane and desflurane groups (Fig. 2A, B). Next, we inspected gene expression levels in the livers of rats subjected to inhalational anesthesia with desflurane as to genes that were contributed to the identification of the following GO terms as sevoflurane-specific terms because of their downregulation more than 2-fold by sevoflurane treatment: “defense response (GO:0006952)”, “response to bacterium (GO:0009617)”, or “defense response to virus (GO:0051607)”. A heatmap visualization of the gene expressions revealed that most genes tended to reduce their expression levels also by desflurane treatment (Fig. S2A). Likewise, most genes responsible for the identification of “positive regulation of transcription from RNA polymerase II promoter (GO:0045944)” or “positive regulation of JNK cascade (GO:0046330)” as desflurane treatment-specific GO terms had a tendency to have reduced expression levels also by sevoflurane treatment (Fig. S2B), indicating that sevoflurane and desflurane exerted similar effects also on transcriptional repression. Consistent with this notion, Venn diagrams (Fig. 2C) showed that significant numbers of genes were commonly repressed by desflurane and sevoflurane (121 genes), although the commonality ratio was not as large as that for the upregulated genes. Given that there was significant similarity between the effects of sevoflurane and desflurane on transcriptional repression and activation, a gene set comprised of the term “cholesterol biosynthetic process (GO:0006695)” appeared to be an exception. Indeed, the expressions of six out of ten genes that contributed to the identification of this term (GO:0006695) as a sevoflurane treatment-specific GO term due to their reduction in expression levels greater than 2-fold by sevoflurane treatment were not significantly altered or even activated by desflurane (Fig. S2C). Therefore, we extended the analyses using all members of genes constituting this GO term. These analyses further clarified the differences in effects on expression of genes related to cholesterol biosynthetic process between sevoflurane and desflurane in which genes downregulated were not restricted to the above ten genes, but vast majority of genes constituting this GO term tended to have reduced expression levels after sevoflurane treatment, whereas most of them did not significantly alter their expression levels after desflurane treatment (Fig. 2D).

**Figure 2.**
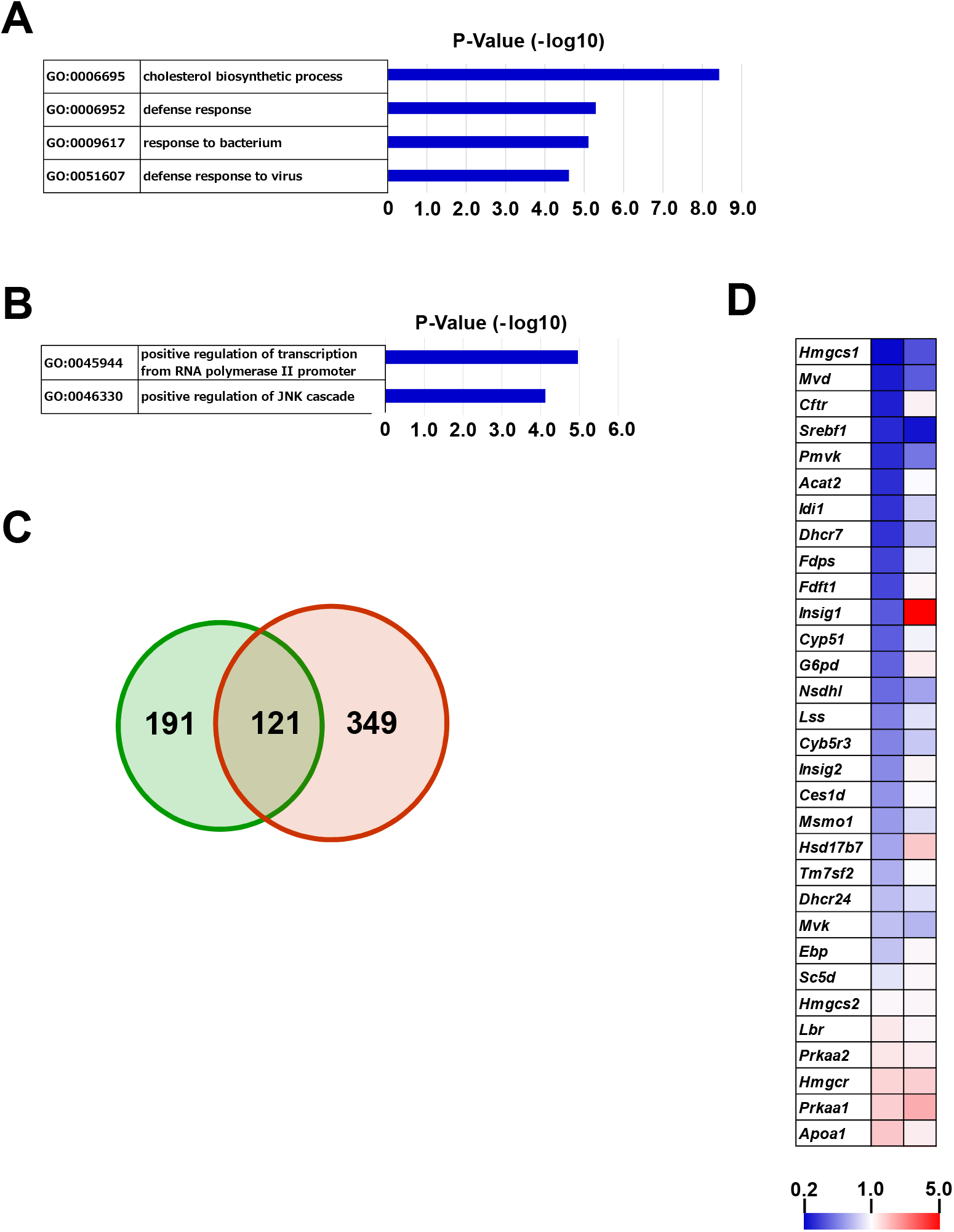
GO analyses of genes downregulated by inhalational anesthesia. Genes whose signal (TPM) values from RNA sequencing decreased more than 2-fold after anesthesia induced by sevoflurane (A) or desflurane (B) were individually subjected to GO analyses as in Figure 1. (C) Venn diagram showing a comparison of genes downregulated more than 2-fold by sevoflurane or desflurane. (D) Heatmap showing a comparison of alterations in expression levels of genes constituting the GO term “cholesterol biosynthetic process (0006695)” by sevoflurane (left column) and desflurane (right column).

### Identification of gene sets coordinately regulated by desflurane and/or sevoflurane via gene set enrichment analysis

In addition to the above GO analyses, we also conducted GSEA to assess similarities and dissimilarities in phenotypic changes that occurred in rat livers subjected to inhalational anesthesia between sevoflurane and desflurane (Fig. S3). In the analyses, we used three publicly available databases, “biological process of Gene Ontology”, “Kyoto Encyclopedia of Genes and Genome”, and “Reactome Pathway Database”. First, we found that gene sets related to circadian rhythm (REGULATION_OF_CIRCADIAN_SLEEP_WAKE_CYCLE; RHYTHMIC_BEHAVIOR) were included as commonly activated gene sets after sevoflurane and desflurane treatment (Fig. 3A). We also found a term related to cholesterol biosynthesis (CHOLESTEROL_BIOSYSNTHESIS) was identified specifically by sevoflurane treatment as a set of genes that were coordinately downregulated (Fig. 3B), validating the data obtained by GO analyses. A new finding of the analyses was a term related to drug metabolism (DRUG_METABOLISM_OTHER_ENZYMES), which was identified as a commonly activated gene set (Fig. 3C). This suggested that desflurane treatment also activated substantial numbers of genes related to drug metabolism, as previously reported for sevoflurane treatment^29,30^.

**Figure 3.**
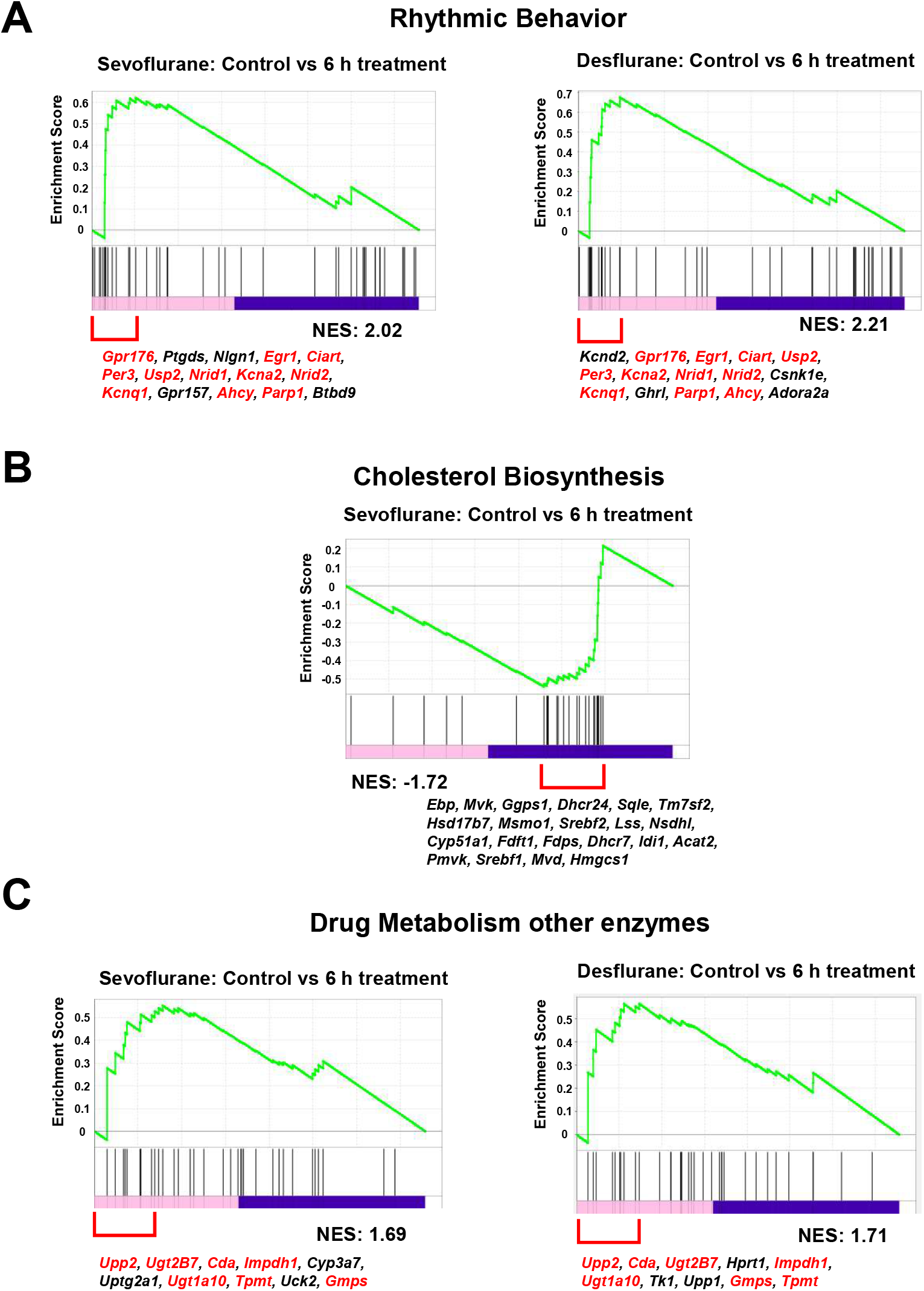
GSEA analyses. (A) Snapshots showing a tendency for the positive regulation of genes constituting the term “Rhythmic Behavior” after treatment with sevoflurane (left panel) or desflurane (right panel). A list of genes denoted as leading-edge genes by the analysis is provided under each snapshot. Genes that are common to both treatments are indicated with red letters. (B) Snapshots showing a tendency for the negative regulation of genes constituting the term “Cholesterol Biosynthesis” after treatment with sevoflurane. A list of leading-edge genes is provided as above. (C) Snapshots showing a tendency for the positive regulation of genes constituting the term “Drug Metabolism Other Enzymes” after treatment with sevoflurane (left panel) or desflurane (right panel). A list of leading-edge genes is provided in which genes specific to either treatment and those commonly identified after both treatments are indicated by black and red letters, respectively, as above.

### Validation of global gene expression analysis data by qPCR analyses of representative genes

Next, we conducted qPCR analyses of genes whose expression levels were significantly up- or downregulated by sevoflurane and/or desflurane by means of global gene expression analyses. Specifically, we selected four (*Alas, Tp53i13, Cyp2b1*, and *Cyp7a1*) (Fig. 4A), two (*Per3* and *Dbp*) (Fig. 4B), and two (*Fdps* and *Fdft1*) genes (Fig. 4C) as representative genes related to “response to cyclic organic compound”, “circadian rhythm”, and “cholesterol biosynthetic process”, respectively, for the analyses. In addition, we examined the expression levels of five additional genes whose expressions were indicated to be markedly upregulated (Fig. S4A) or downregulated (Fig. S4B) by inhalational anesthesia using sevoflurane previously^29^ and/or by our present study using sevoflurane and desflurane. We found that all data from the qPCR analyses were consistent with data from the global gene expression analyses, validating data from both analyses.

**Figure 4.**
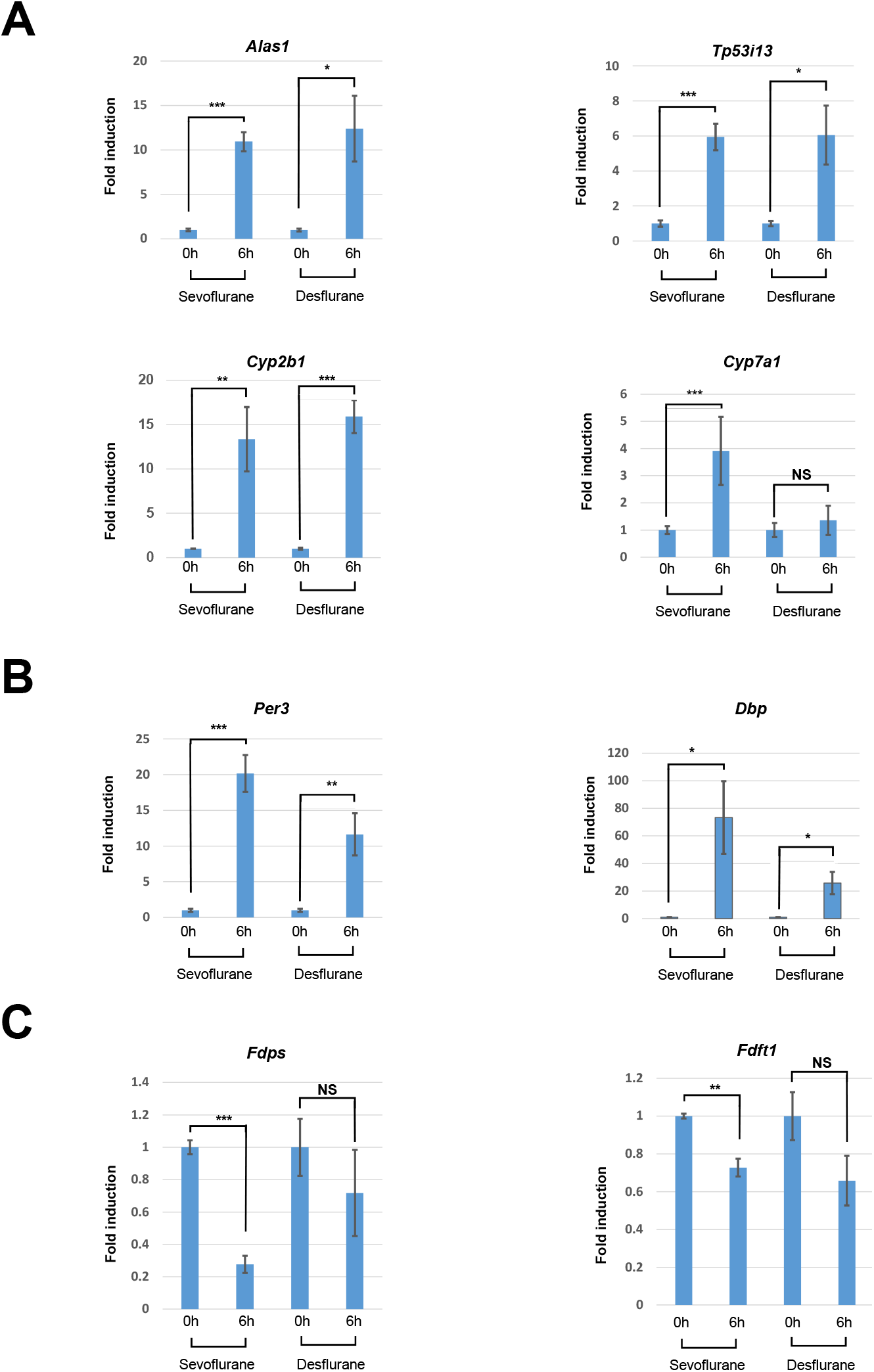
qPCR analyses of representative genes whose expressions were markedly altered by inhalational anesthesia. qPCR analyses of the expressions of four (A), two (B), and two (C) genes as representative members of the gene sets for the terms “response to cyclic organic compound (GO:0014070)”, “circadian rhythm (GO:0032922)”, and “cholesterol biosynthetic process (GO:0006695)”, respectively. Data from control rats in which livers were recovered immediately after the loss of consciousness by sevoflurane or desflurane treatment were arbitrarily set to one. Data represent the mean ± SD of three independent experiments. The Student’s *t*-test was conducted to examine statistical significance. *** *P*<0.001, ** *P*<0.01, * *P*<0.05, NS (not significant) *P*>0.05.

## Discussion

Sevoflurane and desflurane are the most commonly used inhalational anesthetic agents in modern anesthesia practice^33,34^. However, since these two halogenated anesthetics have never been compared with respect to alterations in global expression profiles in livers that may allow to assess the levels of their stress and damage, we conducted next-generation sequence analyses using mRNAs from livers of rats subjected to inhalational anesthesia using sevoflurane or desflurane. Our data revealed that many genes activated and repressed by inhalational anesthesia using sevoflurane were respectively up- and downregulated, indicating that sevoflurane and desflurane caused similar biological effects. Our bioinformatics analyses (GO and GSEA) reinforced these findings, especially for the positively influenced biological effects. Indeed, these analyses revealed that drug metabolism and circadian rhythmic processes that were previously shown to be activated by sevoflurane^29,30^ were also significantly activated by desflurane. Although neither GO nor GSEA analyses indicated any obvious similarities with respect to negatively influenced biological effects, heatmap analyses revealed that many genes that contributed to the identification of sevoflurane treatment-specific terms by their downregulation more than 2-fold also tended to have reduced expression by desflurane. Likewise, many genes that contributed to the identification of desflurane treatment-specific GO terms by the downregulation of their expression levels after desflurane treatment were also downregulated by sevoflurane. Taken together, our data indicate that alterations in the global expression profiles of genes in rat livers, which might correlate with levels of hepatotoxicity, were comparable between the sevoflurane and desflurane groups. However, most genes involved in the cholesterol biosynthetic process had markedly downregulated expression levels after sevoflurane treatment, but not after desflurane treatment. The mechanisms involved in the differential regulation of these genes by sevoflurane and desflurane are not known at present and therefore require further study. In addition, we will investigate the possibility that, unlike the wild-type animals used in this study, sevoflurane and desflurane induce different levels of hepatotoxicity in experimental animals or humans genetically or non-genetically defective in cholesterol biosynthesis.

## Supporting information

Nogi et al. Text and Figures

## Acknowledgments

The authors thank Macrogen, Inc. for RNA-sequencing analyses. We thank J. Ludovic Croxford, PhD, from Edanz (https://jp.edanz.com/ac) for editing a draft of this manuscript.

