## Supplementary material for "Similarity and dissimilarity in alteration of gene expression profile associated with inhalational anesthesia between sevoflurane and desflurane": Nogi et al. Text and Figures

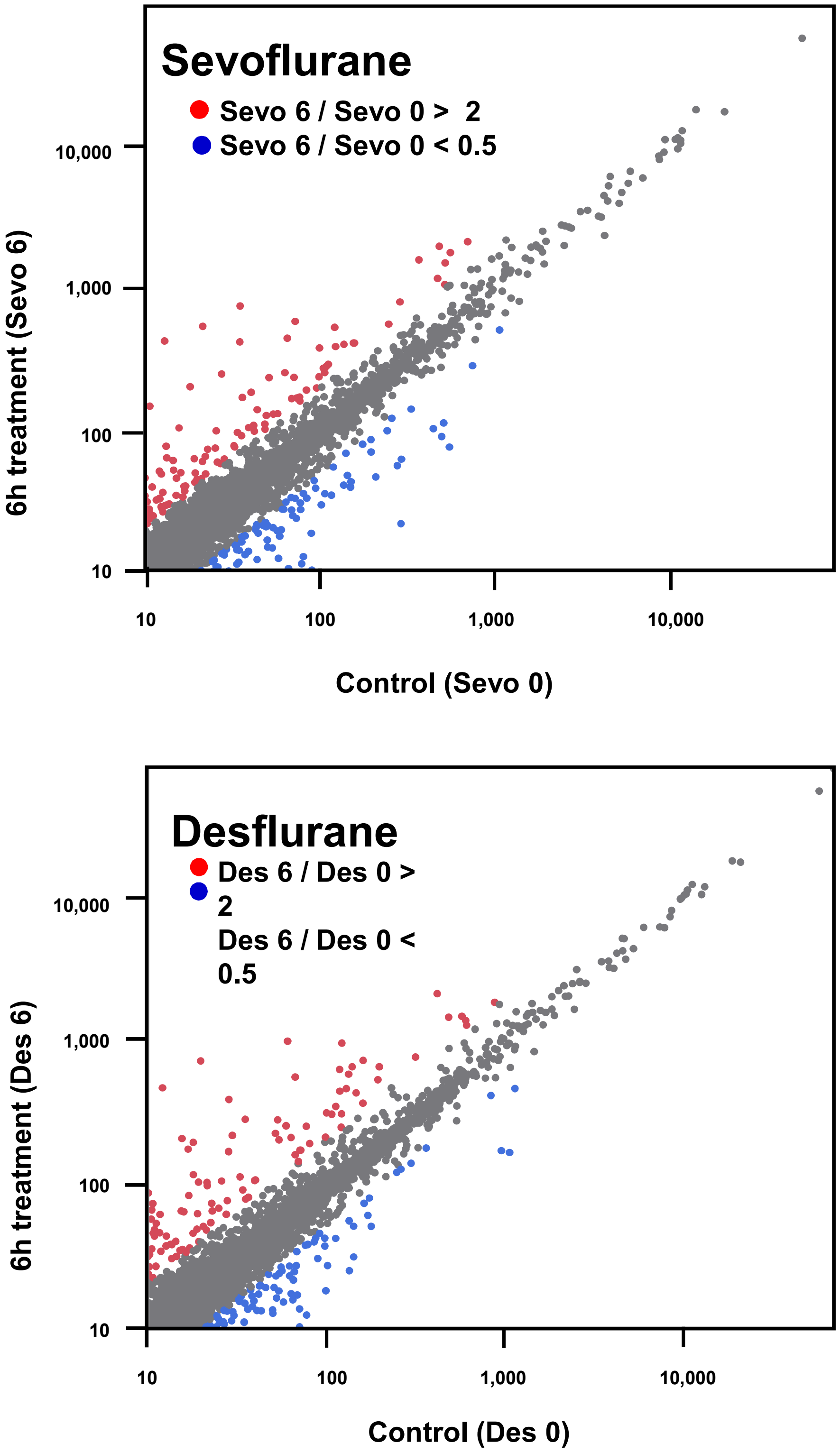


**Figure S1. Scatter plot of RNA sequence data.** Upper and lower panels show data obtained from experiments of inhalational anesthesia using sevoflurane and desflurane, respectively. Numerical values shown on the X- and Y-axes are TPM values from RNA sequence data. Genes whose TPM values were increased or decreased more than 2-fold by inhalational anesthesia using sevoflurane or desflurane are indicated in red and blue, respectively. The numbers of genes up- and downregulated by 6 h treatment with sevoflurane were 434 and 312, respectively, and 350 and 470 genes were up- and downregulated by desflurane treatment, respectively, as shown in Table S1.

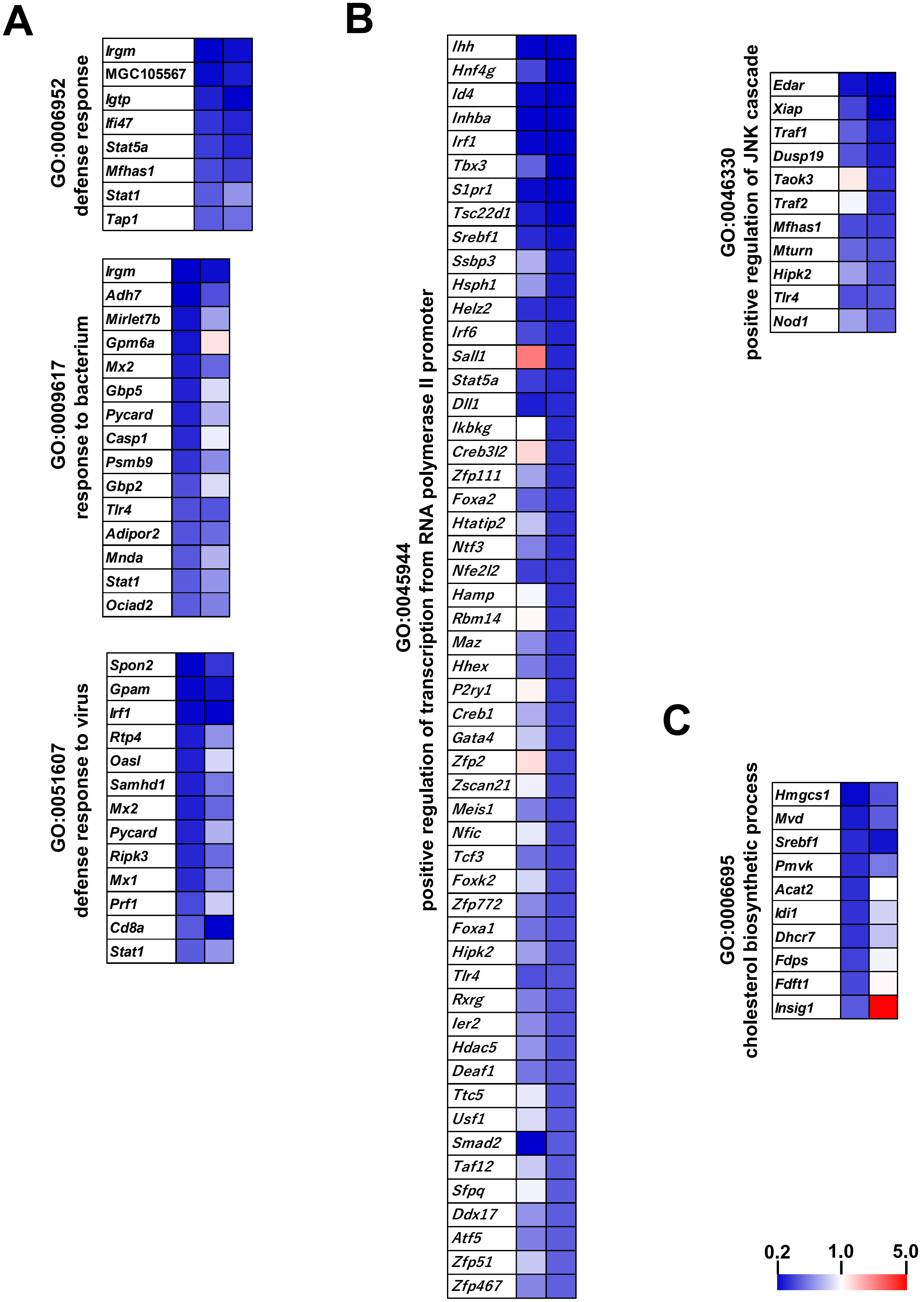
A
GO:0006952
defense response
Irgm
MGC105567
Igtp
Ifi47
Stat5a
Mfhas1
Stat1
Tap1
GO:0009617
response to bacterium
Irgm
Adh7
Mirlet7b
Gpm6a
Mx2
Gbp5
Pycard
Casp1
Psmb9
Gbp2
Tlr4
Adipor2
Mnda
Stat1
Ociad2
GO:0051607
defense response to virus
Spon2
Gpam
Irf1
Rtp4
Oasl
Samhd1
Mx2
Pycard
Ripk3
Mx1
Prf1
Cd8a
Stat1
B
GO:0045944
positive regulation of transcription from RNA polymerase II promoter
Ihh
Hnf4g
Id4
Inhba
Irf1
Tbx3
S1pr1
Tsc22d1
Srebf1
Ssbp3
Hsph1
Helz2
Irf6
Sall1
Stat5a
Dll1
Ikbkg
Creb3l2
Zfp111
Foxa2
Htatip2
Ntf3
Nfe2l2
Hamp
Rbm14
Maz
Hhex
P2ry1
Creb1
Gata4
Zfp2
Zscan21
Meis1
Nfic
Tcf3
Foxk2
Zfp772
Foxa1
Hipk2
Tlr4
Rxrg
Ier2
Hdac5
Deaf1
Ttc5
Usf1
Smad2
Taf12
Sfpq
Ddx17
Atf5
Zfp51
Zfp467
GO:0046330
positive regulation of JNK cascade
Edar
Xiap
Traf1
Dusp19
Taok3
Traf2
Mfhas1
Mturn
Hipk2
Tlr4
Nod1
C
GO:0006695
cholesterol biosynthetic process
Hmgcs1
Mvd
Srebf1
Pmvk
Acat2
Idi1
Dhcr7
Fdps
Fdft1
Insig1
0.2
1.0
5.0

**Figure S2. Heatmap comparisons of genes downregulated by inhalational anesthesia with sevoflurane or desflurane.** (A) Effect of desflurane treatment on the expressions of genes that contribute to the identification of sevoflurane treatment-specific GO terms. Genes downregulated more than 2-fold by sevoflurane treatment were selected among genes constituting the GO terms of “defense response (0006952)”, “response to bacterium (0009617)”, or “defense response to virus (051607)”. Their expression levels in the livers of rats treated with desflurane for 6 h were demonstrated by a heatmap (right column) along with data obtained by the analyses of livers of rats treated with sevoflurane (left column). (B) Effect of sevoflurane treatment on the expressions of genes that contributed to the identification of desflurane treatment-specific GO terms. Genes downregulated more than 2-fold by desflurane treatment were selected among genes constituting the GO terms “positive regulation of transcription from RNA polymerase II promoter (0045944)” or “positive regulation of JNK cascade (0046330)”. Their expression levels in livers of rats treated with desflurane (right column) or sevoflurane (left column) for 6 h were demonstrated by a heatmap as in (A). (C) Effect of desflurane treatment on the expressions of genes that contributed to the identification of “cholesterol biosynthetic process (0006695)” as a sevoflurane treatment-specific GO term. Genes downregulated more than 2-fold by sevoflurane treatment were selected among genes constituting the GO term “cholesterol biosynthetic process (0006695)” and their expression levels in livers of rats treated with desflurane (right column) or sevoflurane (left column) for 6 h were demonstrated by a heatmap as in (A).

A

#### GOBP

| Sevoflurane 0h < 6h | NES | P-value | Desflurane 0h < 6h | NES | P-value |
|---|---|---|---|---|---|
| SODIUM_ION_HOMEOSTASIS | 2.33 | 0 | NEGATIVE_REGULATION_OF_VASCULAR_ENDOTHELIAL_GROWTH_FACTOR_PRODUCTION | 2.32 | 0 |
| MIRNA_MEDIATED_GENE_SILENCING_BY_INHIBITION_OF_TRANSLATION | 2.23 | 0 | RHYTHMIC_BEHAVIOR | 2.21 | 0 |
| GLIAL_CELL_PROLIFERATION | 2.05 | 0 | NEGATIVE_REGULATION_OF_AMYLOID_PRECURSOR_PROTEIN_BIOSYNTHETIC_PROCESS | 2.16 | 0 |
| CELLULAR_SODIUM_ION_HOMEOSTASIS | 2.05 | 0.001 | POSITIVE_REGULATION_OF_NUCLEOTIDE_BIOSYNTHETIC_PROCESS | 2.13 | 0.001 |
| NEGATIVE_REGULATION_OF_VASCULAR_ENDOTHELIAL_GROWTH_FACTOR_PRODUCTION | 2.03 | 0.003 | POSITIVE_REGULATION_OF_GLUCONEOGENESIS | 2.11 | 0 |
| RHYTHMIC_BEHAVIOR | 2.02 | 0 | RESPONSE_TO_DIETARY_EXCESS | 2.1 | 0.001 |
| ACROSOME_REACTION | 2.02 | 0.004 | AMYLOID_PRECURSOR_PROTEIN_BIOSYNTHETIC_PROCESS | 2.1 | 0 |
| REGULATION_OF_CIRCADIAN_SLEEP_WAKE_CYCLE | 2.01 | 0.001 | EATING_BEHAVIOR | 2.09 | 0 |
| POSITIVE_REGULATION_OF_GLIAL_CELL_PROLIFERATION | 2.01 | 0.004 | NEGATIVE_REGULATION_OF_AMYLOID_PRECURSOR_PROTEIN_CATABOLIC_PROCESS | 2.08 | 0 |
| POSITIVE_REGULATION_OF_GLUCONEOGENESIS | 2.01 | 0.001 | REGULATION_OF_NUCLEOTIDE_BIOSYNTHETIC_PROCESS | 2.07 | 0 |
| REGULATION_OF_GLIAL_CELL_PROLIFERATION | 1.97 | 0.005 | MIRNA_MEDIATED_GENE_SILENCING_BY_INHIBITION_OF_TRANSLATION | 2.06 | 0 |
| ENERGY_HOMEOSTASIS | 1.97 | 0.001 | MONOVALENT_INORGANIC_ANION_HOMEOSTASIS | 2.06 | 0 |
| MIDBRAIN_DOPAMINERGIC_NEURON_DIFFERENTIATION | 1.97 | 0.004 | NEUTRAL_LIPID_BIOSYNTHETIC_PROCESS | 2.06 | 0 |
| CATECHOLAMINE_UPTAKE | 1.96 | 0.001 | PLASMA_MEMBRANE_FUSION | 2.05 | 0 |
| POSITIVE_REGULATION_OF_VASCULAR_ASSOCIATED_SMOOTH_MUSCLE_CELL_MIGRATION | 1.95 | 0.008 | POSITIVE_REGULATION_OF_EXTRACELLULAR_MATRIX_ORGANIZATION | 2.05 | 0 |
| TRABECULA_FORMATION | 1.95 | 0.003 | FUSION_OF_SPERM_TO_EGG_PLASMA_MEMBRANE_INVOLVED_IN_SINGLE_FERTILIZATION | 2.03 | 0 |
| CIRCADIAN_SLEEP_WAKE_CYCLE | 1.95 | 0.004 | TRABECULA_FORMATION | 2.03 | 0.004 |
| MITOCHONDRIAL_GENOME_MAINTENANCE | 1.92 | 0.006 | DETECTION_OF_MECHANICAL_STIMULUS | 2.01 | 0 |
| PYRIMIDINE_CONTAINING_COMPOUND_CATABOLIC_PROCESS | 1.91 | 0.005 | THIOESTER_BIOSYNTHETIC_PROCESS | 2 | 0.002 |
| REGULATION_OF_CILIUM_DEPENDENT_CELL_MOTILITY | 1.89 | 0.009 | POSITIVE_REGULATION_OF_CYCLIN_DEPENDENT_PROTEIN_KINASE_ACTIVITY | 1.98 | 0.006 |

#### KEGG

| Sevoflurane | NES | P-value | Desflurane | NES | P-value |
|---|---|---|---|---|---|
| DRUG_METABOLISM_OTHER_ENZYMES | 1.69 | 0.024 | OLFACTORY_TRANSDUCTION | 2.13 | 0 |
| ANTIGEN_PROCESSING_AND_PRESENTATION | 1.68 | 0.036 | ASTHMA | 1.95 | 0.003 |
| PROXIMAL_TUBULE_BICARBONATE_RECLAMATION | 1.68 | 0.047 | ECM_RECEPTOR_INTERACTION | 1.91 | 0.006 |
| | | | COMPLEMENT_AND_COAGULATION_CASCADES | 1.86 | 0.005 |
| | | | ERBB_SIGNALING_PATHWAY | 1.81 | 0.009 |
| | | | CITRATE_CYCLE_TCA_CYCLE | 1.78 | 0.014 |
| | | | LINOLEIC_ACID_METABOLISM | 1.73 | 0.02 |
| | | | DRUG_METABOLISM_OTHER_ENZYMES | 1.71 | 0.026 |
| | | | ALLOGRAFT_REJECTION | 1.69 | 0.031 |
| | | | LEISHMANIA_INFECTION | 1.61 | 0.03 |
| | | | NATURAL_KILLER_CELL_MEDIATED_CYTOTOXICITY | 1.58 | 0.024 |
| | | | ADIPOCYTOKINE_SIGNALING_PATHWAY | 1.58 | 0.035 |
| | | | GNRH_SIGNALING_PATHWAY | 1.56 | 0.035 |
| | | | NEUROTROPHIN_SIGNALING_PATHWAY | 1.53 | 0.033 |

#### REACTOME

| Sevoflurane | NES | P-value | Desflurane | NES | P-value |
|---|---|---|---|---|---|
| DIGESTION_AND_ABSORPTION | 2.26 | 0 | DIGESTION_AND_ABSORPTION | 2.19 | 0 |
| WNT_LIGAND_BIOGENESIS_AND_TRAFFICKING | 2.11 | 0.003 | COLLAGEN_BIOSYNTHESIS_AND_MODIFYING_ENZYMES | 2.15 | 0 |
| DIGESTION | 2.1 | 0.001 | FORMATION_OF_FIBRIN_CLOT_CLOTTING_CASCADE | 2.1 | 0 |
| NEGATIVE_REGULATION_OF_TCF_DEPENDENT_SIGNALING_BY_WNT_LIGAND_ANTAGONISTS | 1.93 | 0.003 | COLLAGEN_CHAIN_TRIMERIZATION | 2.1 | 0.004 |
| SYNAPTIC_ADHESION_LIKE_MOLECULES | 1.9 | 0.009 | COLLAGEN_FORMATION | 2.05 | 0.002 |
| DOPAMINE_NEUROTRANSMITTER_RELEASE_CYCLE | 1.89 | 0.011 | INTRINSIC_PATHWAY_OF_FIBRIN_CLOT_FORMATION | 2.04 | 0.001 |
| NEUROTRANSMITTER_RELEASE_CYCLE | 1.86 | 0.011 | ECM_PROTEOGLYCANS | 2.01 | 0.001 |
| SEROTONIN_NEUROTRANSMITTER_RELEASE_CYCLE | 1.85 | 0.008 | NCAM1_INTERACTIONS | 1.99 | 0.001 |
| RRNA_MODIFICATION_IN_THE_NUCLEUS_AND_CYTOSOL | 1.84 | 0.008 | CROSSLINKING_OF_COLLAGEN_FIBRILS | 1.99 | 0 |
| PHASE_2_PLATEAU_PHASE | 1.84 | 0.007 | TRANSPORT_OF_INORGANIC_CATIONS_ANIONS_AND_AMINO_ACIDS_OLIGOPEPTIDES | 1.96 | 0.001 |
| GLYCOGEN_METABOLISM | 1.84 | 0.013 | ASSEMBLY_OF_COLLAGEN_FIBRILS_AND_OTHER_MULTIMERIC_STRUCTURES | 1.93 | 0.004 |
| CIRCADIAN_CLOCK | 1.79 | 0.011 | NCAM_SIGNALING_FOR_NEURITE_OUT_GROWTH | 1.91 | 0.007 |
| N_GLYCAN_ANTENNAE_ELONGATION | 1.78 | 0.014 | AMINO_ACID_TRANSPORT_ACROSS_THE_PLASMA_MEMBRANE | 1.9 | 0.003 |
| TRANSPORT_OF_BILE_SALTS_AND_ORGANIC_ACIDS_METAL_IONS_AND_AMINE_COMPOUNDS | 1.74 | 0.011 | NON_INTEGRIN_MEMBRANE_ECM_INTERACTIONS | 1.87 | 0 |
| DISEASES_OF_DNA_REPAIR | 1.74 | 0.015 | DEFECTS_OF_CONTACT_ACTIVATION_SYSTEM_CAS_AND_KALLIKREIN_KININ_SYSTEM_KKS | 1.87 | 0.011 |
| HOMOLOGOUS_DNA_PAIRING_AND_STRAND_EXCHANGE | 1.67 | 0.037 | NGF_STIMULATED_TRANSCRIPTION | 1.87 | 0.007 |
| GLYCOGEN_STORAGE_DISEASES | 1.65 | 0.04 | DEFECTIVE_INTRINSIC_PATHWAY_FOR_APOPTOSIS | 1.84 | 0.007 |
| RNA_POLYMERASE_I_TRANSCRIPTION_TERMINATION | 1.65 | 0.032 | SCAVENGING_BY_CLASS_A_RECEPTORS | 1.84 | 0.008 |
| UNFOLDED_PROTEIN_RESPONSE_UPR | 1.64 | 0.027 | SLC_MEDIATED_TRANSMEMBRANE_TRANSPORT | 1.84 | 0 |
| NON_INTEGRIN_MEMBRANE_ECM_INTERACTIONS | 1.63 | 0.032 | INTERACTION_BETWEEN_L1_AND_ANKYRINS | 1.83 | 0.013 |

## B

##### GOBP

| Sevoflurane 0h > 6h | NES | P-value | Desflurane 0h > 6h | NES | P-value |
|---|---|---|---|---|---|
| NEUROPEPTIDE_SIGNALING_PATHWAY | -1.93 | 0 | MYOTUBE_CELL_DEVELOPMENT | -2.06 | 0 |
| NEUROMUSCULAR_SYNAPTIC_TRANSMISSION | -1.927 | 0.0048544 | SKELETAL_MUSCLE_ADAPTATION | -2.03 | 0 |
| NATURAL_KILLER_CELL_ACTIVATION_INVOLVED_IN_IMMUNE_RESPONSE | -1.883 | 0 | ACTIN_MYOSIN_FILAMENT_SLIDING | -2.01 | 0 |
| CARDIAC_MYOFIBRIL_ASSEMBLY | -1.85 | 0 | CARDIAC_CELL_FATE_COMMITMENT | -1.86 | 0 |
| MONOCYTE_CHEMOTACTIC_PROTEIN_1_PRODUCTION | -1.823 | 0.0155642 | ATRIAL_CARDIAC_MUSCLE_TISSUE_DEVELOPMENT | -1.86 | 0.012 |
| POSITIVE_REGULATION_OF_CIRCADIAN_RHYTHM | -1.816 | 0 | POSITIVE_REGULATION_OF_STRIATED_MUSCLE_CELL_DIFFERENTIATION | -1.85 | 0 |
| VESICLE_FUSION_TO_PLASMA_MEMBRANE | -1.813 | 0 | PIRNA_METABOLIC_PROCESS | -1.8 | 0.003 |
| NEGATIVE_REGULATION_OF_LYASE_ACTIVITY | -1.811 | 0 | CALCIUM_ION_REGULATED_EXOCYTOSIS_OF_NEUROTRANSMITTER | -1.78 | 0 |
| POSITIVE_REGULATION_OF_INTERLEUKIN_12_PRODUCTION | -1.788 | 0 | EMBRYONIC_SKELETAL_JOINT_DEVELOPMENT | -1.78 | 0.003 |
| POSITIVE_REGULATION_OF_MONOCYTE_CHEMOTACTIC_PROTEIN_1_PRODUCTION | -1.786 | 0.0037453 | REGULATION_OF_ODONTOGENESIS | -1.75 | 0.004 |
| NEGATIVE_REGULATION_OF_ADENYLATE_CYCLASE_ACTIVITY | -1.78 | 0.0042017 | REGULATION_OF_CARDIAC_MUSCLE_CELL_DIFFERENTIATION | -1.75 | 0.012 |
| GONADOTROPIN_SECRETION | -1.74 | 0.0068259 | ANATOMICAL_STRUCTURE_REGRESSION | -1.74 | 0.01 |
| ANTIMICROBIAL_HUMORAL_RESPONSE | -1.702 | 0 | REGULATION_OF_CARDIOCYTE_DIFFERENTIATION | -1.71 | 0.015 |
| CELLULAR_RESPONSE_TO_INTERFERON_BETA | -1.686 | 0.0082305 | KERATINIZATION | -1.69 | 0.011 |
| CALCIUM_ION_TRANSPORT_INTO_CYTOSOL | -1.685 | 0.0147059 | REGULATION_OF_MYOBLAST_PROLIFERATION | -1.68 | 0.004 |
| NITRIC_OXIDE_MEDIATED_SIGNAL_TRANSDUCTION | -1.666 | 0.0206612 | POSITIVE_REGULATION_OF_MYOTUBE_DIFFERENTIATION | -1.67 | 0.003 |
| RESPONSE_TO_SALT_STRESS | -1.665 | 0.0047847 | POSITIVE_REGULATION_OF_TISSUE_REMODELING | -1.66 | 0.008 |
| EMBRYONIC_SKELETAL_JOINT_DEVELOPMENT | -1.663 | 0.0117188 | SULFATION | -1.64 | 0.013 |
| REGULATION_OF_INOSITOL_PHOSPHATE_BIOSYNTHETIC_PROCESS | -1.651 | 0.0114504 | REGULATION_OF_B_CELL_DIFFERENTIATION | -1.64 | 0.02 |
| ACTIVATION_OF_PHOSPHOLIPASE_C_ACTIVITY | -1.649 | 0.015625 | RESPONSE_TO_FUNGUS | -1.64 | 0.014 |

##### KEGG

| Sevoflurane 0h > 6h | NES | P-value | Desflurane 0h > 6h | NES | P-value |
|---|---|---|---|---|---|
| STEROID_BIOSYNTHESIS | -1.54 | 0.042 | SNARE_INTERACTIONS_IN_VESICULAR_TRANSPORT | -1.64 | 0.014 |
| RIBOSOME | -1.42 | 0 | MATURITY_ONSET_DIABETES_OF_THE_YOUNG | -1.5 | 0.035 |
| NOD_LIKE_RECEPTOR_SIGNALING_PATHWAY | -1.3 | 0.082 | PRIMARY_IMMUNODEFICIENCY | -1.5 | 0.036 |
| OLFACTORY_TRANSDUCTION | -1.27 | 0 | | | |

##### REACTOME

| Sevoflurane 0h > 6h | NES | P-value | Desflurane 0h > 6h | NES | P-value |
|---|---|---|---|---|---|
| DEFENSINS | -2.18 | 0 | CLASS_C_3_METABOTROPIC_GLUTAMATE_PHEROMONE_RECEPTORS | -1.9524 | 0 |
| BETA_DEFENSINS | -2.06 | 0 | CHEMOKINE_RECEPTORS_BIND_CHEMOKINES | -1.9176 | 0 |
| METABOLISM_OF_AMINE_DERIVED_HORMONES | -1.96 | 0 | CYTOSOLIC_SULFONATION_OF_SMALL_MOLECULES | -1.7184 | 0.0037879 |
| CHOLESTEROL_BIOSYNTHESIS | -1.72 | 0.01 | STRIATED_MUSCLE_CONTRACTION | -1.7097 | 0.0050761 |
| FORMATION_OF_TUBULIN_FOLDING_INTERMEDIATES_BY_CCT_TRIC | -1.69 | 0.013 | BETA_DEFENSINS | -1.6351 | 0.0221239 |
| RHO_GTPASES_ACTIVATE_IQGAPS | -1.66 | 0.014 | GERM_LAYER_FORMATION_AT_GASTRULATION | -1.6256 | 0.0338983 |
| EUKARYOTIC_TRANSLATION_ELONGATION | -1.62 | 0 | SENSORY_PERCEPTION_OF_TASTE | -1.5689 | 0.0310881 |
| POST_CHAPERONIN_TUBULIN_FOLDING_PATHWAY | -1.59 | 0.036 | CLASS_B_2_SECRETIN_FAMILY_RECEPTORS | -1.5628 | 0 |
| INTERFERON_ALPHA_BETA_SIGNALING | -1.58 | 0.024 | GLUCAGON_TYPE_LIGAND_RECEPTORS | -1.4951 | 0.0380952 |
| AMINE_LIGAND_BINDING_RECEPTORS | -1.54 | 0.02 | NEUREXINS_AND_NEUROLIGINS | -1.4404 | 0.0440252 |
| ANTIMICROBIAL_PEPTIDES | -1.54 | 0.012 | | | |
| IKK_COMPLEX_RECRUITMENT_MEDIATED_BY_RIP1 | -1.48 | 0.037 | | | |
| ACTIVATION_OF_GENE_EXPRESSION_BY_SREBF_SREBP | -1.48 | 0.021 | | | |
| COOPERATION_OF_PREFOLDIN_AND_TRIC_CCT_IN_ACTIN_AND_TUBU | -1.47 | 0.046 | | | |
| INWARDLY_RECTIFYING_K_CHANNELS | -1.47 | 0.047 | | | |
| GABA_RECEPTOR_ACTIVATION | -1.45 | 0.036 | | | |
| AGGREPHAGY | -1.43 | 0.041 | | | |
| PEPTIDE_LIGAND_BINDING_RECEPTORS | -1.43 | 0 | | | |

**Figure S3. GSEA analyses of RNA sequence data from livers of rats subjected to inhalational anesthesia.** (A) A list of gene sets identified by GSEA analyses as significantly activated gene sets by inhalational anesthesia using sevoflurane or desflurane. Three distinct publicly available databases, “biological process of Gene Ontology”, “Kyoto Encyclopedia of Genes and Genome”, and “Reactome Pathway Database” were used for the analyses in which the top twenty terms according to their values of normalized enrichment score were selected from positively-regulated gene sets with the analyses using each platform, but terms whose *P*-values were greater than 0.05 were eliminated from the list. Commonly identified gene sets by treatment with sevoflurane or desflurane were individually marked by distinct colors. (B) A list of gene sets identified by GSEA analyses as significantly repressed gene sets by inhalational anesthesia using sevoflurane or desflurane. The same criteria used in (A) were used and the terms of the gene set that met the criteria are shown. Commonly identified gene sets by treatment with sevoflurane or desflurane were individually marked by light brown or dull blue.

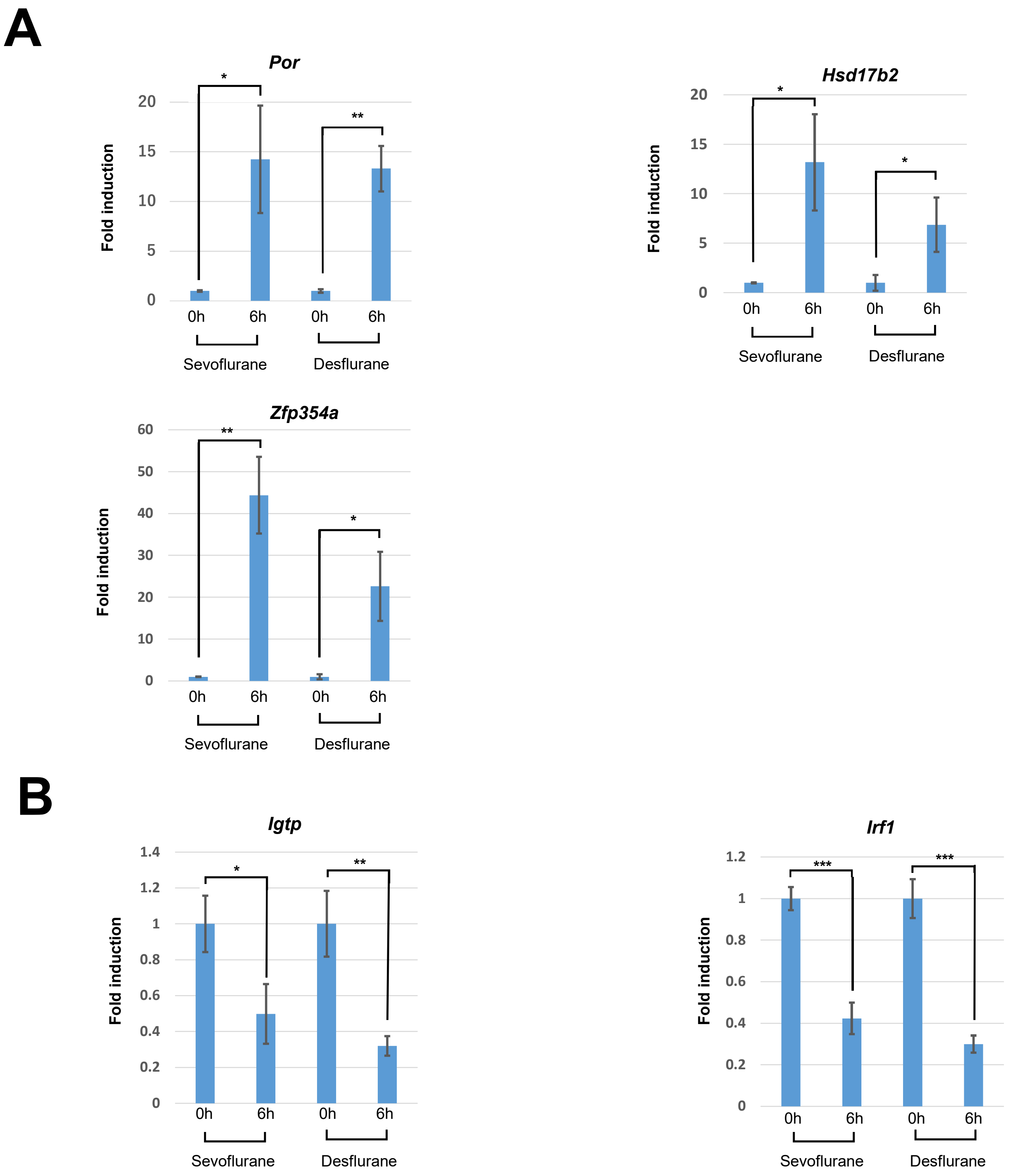


**Figure S4. qPCR analyses of genes whose expressions were markedly altered by inhalational anesthesia in this study and/or previous studies.** (A) qPCR to quantify the expression levels of genes markedly activated by previous DNA microarray[29] and RNA sequence analyses in the current study. Data from control rats in which livers were recovered immediately after the loss of rat consciousness induced by sevoflurane or desflurane treatment were arbitrarily set to one. Data represent the mean ± SD of three independent experiments. Statistical analyses were performed as in Figure 4. (B) qPCR to quantify the expression levels of immunity-related genes strongly repressed by a DNA microarray reported by Sakamoto et al.[29] as well as RNA sequence analyses from the current study. Statistical analyses were performed as in (A).

**Table S1**

### Gene lists whose expression levels were up- or down-regulated by sevoflurane and/or desflurane

| Up-regulated | | | Down-regulated | | |
|---|---|---|---|---|---|
| Group 1* | Group 2 | Group 3 | Group 4 | Group 5 | Group 6 |
| (204 genes) | (230 genes) | (146 genes) | (121 genes) | (191 genes) | (349 genes) |
| Snora58 | Mir292 | Mir19a | Arntl | Irs3 | Cxcl1 |
| Ciart | Mir295-1 | Gdf15 | Ppp2r2b | Gpx2 | Ihh |
| Maff | Mir295-2 | Slc16a6 | Gck | St8sia1 | Scrn1 |
| Upp2 | Epm2a | Spsb1 | Gimd1 | Fgf21 | Mir290 |
| Igfbp1 | Mir293 | Npy | Stac3 | LOC689065 | Trim47 |
| Gadd45b | LOC684871 | Pvr | Dipk2a | LOC100363193 | Mir292 |
| Cyp2b1 | Mir291b | Fbxo15 | Chka | Rasl11a | Inhba |
| Sds | Plppr1 | Mir345 | Arhgef19 | Tmem79 | RGD1559600 |
| Rgs16 | Mcm10 | F3 | Slc34a2 | Atp2b2 | Inhbe |
| Lpin1 | Znrd1as1 | Pdlim2 | Osgin1 | Gimap7 | Myorg |
| Dbp | Tsku | Serpine1 | Spon2 | RGD1309808 | Efnb2 |
| Ppargc1a | Cxcr4 | Dusp1 | Srms | Dcakd | Tmod1 |
| Ccne1 | Irak1bp1 | Mir7a1 | Ptk6 | Mir19a | Ltc4s |
| Usp2 | Mir290 | Irs2 | Cyp4a2 | Cth | Ackr3 |
| Rell1 | Chmp4c | Trib3 | Id2 | Bmp4 | Tbx3 |
| Alas1 | Ces2e | Mir6326 | Tubb2a | RGD1305464 | Acyp1 |
| Noct | Lenep | Cebpb | Angptl8 | Adh1 | Rcan1 |
| Slc25a29 | Tob2 | Synpo | Tubb2b | Arl4d | Ccl4 |
| Por | Dusp23 | Klf11 | Irgm | Cidea | Dusp6 |
| Got1 | Il6r | Insig1 | Cyp4a3 | Clec4a | L3hypdh |
| Plk3 | Mir343 | Gzmk | Sebox | Rcan2 | Efna5 |
| Coq10b | Aph1b | Rgs1 | Adh7 | Lonrf3 | RGD1562844 |
| Per3 | Cyp7a1 | Wnt4 | Smad2 | Arhgap8 | Rgs3 |
| Trim36 | Entpd7 | Gpr31 | G0s2 | Rpl39l | B4galt5 |
| Zfp354a | Cyp17a1 | Marchf1 | Mid1ip1 | Mirlet7b | Tppp |
| Ppargc1b | Wsb1 | Alpl | Hsd3b5 | Sgk1 | Cish |
| Slc25a32 | Jade1 | Atf3 | S1pr2 | Iqcg | Macir |
| Retreg1 | Ccdc125 | Slc1a2 | Gpam | Phgdh | Ccnd2 |
| Edem1 | Nhlrc1 | Btg2 | Irf1 | Gpm6a | S100a9 |
| Mfsd2a | Zcchc10 | Pfkfb3 | Efna1 | Cep63 | Chordc1 |
| Eif4ebp3 | Sdr42e1 | Brap | Id4 | Clec4a1 | Fahd1 |
| Mt2A | Tns2 | Zfp37 | Hmgcs1 | Litaf | Kbtbd8 |
| Zfp36 | Slc22a5 | LOC100909539 | MGC105567 | Coq8a | Zfp978 |
| Pptc7 | Rbp7 | Gstm3l | S1pr1 | Gdf15 | Thnsl1 |
| Ppp1r15a | Mir291a | Urb2 | Slc6a6 | Rtp4 | Kank2 |
| Rhbdd2 | Tma16 | Rhob | Pcsk9 | Oasl | Rad51d |
| Cyp2b2 | Mir122 | Unc5b | RGD1561157 | Lrtm2 | Banp |
| Per1 | Alas2 | Zc3h12a | Dnmt3b | Mk1 | Ston2 |
| Ppl | Ahr | Ctdspl | Xcl1 | Sult2b1 | Tnfrsf11b |
| Prr15 | Hacl1 | Btg3 | Sgk2 | Samhd1 | Stbd1 |
| Slc38a2 | Lepr | Dbt | Serpina7 | Mx2 | Pdik1l |
| Ptp4a1 | Polr2i | G6pc | Edar | Gbp5 | Mterf1 |
| Pla2g12a | Pigu | Hnrnpd | Tjp3 | Pycard | Bcl6 |
| RGD1563888 | Rbm19 | Mir186 | Rtp3 | Lnx2 | Rcbtb1 |
| Sik1 | Sall1 | Cdkn1a | Layn | Rab3d | Cdc42ep3 |
| Mafb | Tsc22d3 | Tob1 | Arrdc3 | Cytip | Traf1 |
| Slc25a33 | Tom1l2 | Hivep1 | RGD1565616 | Ripk3 | Rnf144b |
| Jun | Nek6 | Qtrt1 | Jade3 | Casp1 | Cyb5d2 |
| Tp53i13 | Ccnd1 | Acot1 | Cldn2 | Tnfrsf1b | S100a4 |

*** Group 1: Genes activated commonly by sevoflurane and desflurane**
**Group 2: Genes activated specifically by sevoflurane**
**Group 3: Genes activated specifically by desflurane**
**Group 4: Genes represed commonly by sevoflurane and desflurane**
**Group 5: Genes repressed specifically by sevoflurane**
**Group 6: Genes repressed specifically by desflurane**

| | | | | | |
|---|---|---|---|---|---|
| N4bp2l1 | Pcgf1 | Mafk | Kctd7 | Sntb1 | Ssbp3 |
| Fbxo31 | Lama4 | Hopx | Socs2 | Angptl4 | Mcc |
| Map3k5 | Hbb-bs | Poc1a | Mvd | Ddah1 | Smim11a |
| Gys2 | Mir21 | Mknk2 | Dll1 | Etv6 | Kcnj8 |
| Adcy10 | S100a8 | Ppif | Tsc22d1 | Mx1 | Hsph1 |
| RGD1562310 | Irak3 | Thrsp | Ppp1r3b | Mirlet7f-1 | LOC100125364 |
| Klf9 | Abhd13 | Mir3557 | Sigmar1 | Nfkbie | Znrd1as1 |
| Tat | Utp14a | H1f0 | RGD1564865 | Pld1 | Ttf2 |
| Pnpla2 | Nr2c1 | Hnrnpdl | Rilp | Mirlet7f1 | Prkd3 |
| Tdo2 | Ccdc86 | Meox2 | Gfod2 | Pmvk | Prrg2 |
| Ablim3 | Palm2 | Zbtb39 | Igtp | Tmem150c | Kank1 |
| Tp53inp1 | Rgs5 | Fmo2 | Mme | Acsm5 | Tubd1 |
| Ifrd1 | Tanc1 | Cdc37l1 | Tmem140 | Acat2 | Mmp15 |
| Dcaf13 | Pck1 | Cep63 | Lpar6 | Il7 | Zbtb26 |
| Cblb | Slc16a12 | Lmcd1 | Srebf1 | Nkg7 | Klhl7 |
| Mt1 | RT1-S2 | Gpd1 | Aqp8 | Fam89a | Ccdc146 |
| Fkbp5 | Rsl1d1l1 | Csnk1e | Tuba4a | Psmb9 | Ankrd37 |
| Gja4 | Pmm1 | Junb | Klrk1 | Clpx | Abtb2 |
| Pim3 | Mir294 | Efemp2 | Mpi | Idi1 | Sall1 |
| Csrnp1 | Nsun5 | Rplp2 | Dnajc12 | Dhcr7 | Akap5 |
| Slc25a47 | Hbb | Mir101-2 | Slc13a4 | Pgpep1 | Med9 |
| Pdp2 | Fbxo32 | Mknk1 | Slc19a2 | Mthfd2 | Ogg1 |
| Mid1 | Hbb-b1 | Gstm5 | Apol9a | Pbsn | Mab21l3 |
| Slc25a25 | Ptpn21 | Abl1 | Helz2 | Gstm5 | Avpr1a |
| Hsd17b2 | Peg3 | Metrnl | Rtn4rl1 | Afg1l | Uqcc3 |
| Ppm1k | Rapgef4 | Insc | Slc17a9 | Cdh1 | Ptgfr |
| Bcl2l11 | Med25 | Gpatch4 | Ifi47 | Prnp | Cdk2ap2 |
| Map1lc3b | Abcg8 | Mef2d | Hao2 | Prelp | Ikbkg |
| Il1r1 | Qsox2 | Otulin | Bcl2a1 | RGD1309104 | Emd |
| Bcar3 | Nlrp12 | Gnat2 | Wdr91 | Spink3 | Cxcl11 |
| Rab30 | Hba-a2 | Jund | Anks4b | Armcx1 | Creb3l2 |
| Trib1 | Slc25a40 | Ass1 | Notch2 | Ly86 | Snx12 |
| Hps4 | Slc2a2 | Vgll4 | Zfp697 | AA926063 | Ncoa5 |
| Pxmp4 | Hba1 | Ttll9 | R3hdm2 | Ypel2 | Cldn14 |
| Rbm3 | Atl2 | Acy3 | Stat5a | Ppard | Zfp111 |
| Sgms1 | Amn1 | Znhit6 | Nfe2l2 | Wdr82 | Acvr1c |
| Tns1 | Pfkfb1 | Sgsm2 | Fkbp4 | Midn | Fbxl19 |
| Tmem86a | Cyp4b1 | Desi2 | Itga5 | Acsl3 | Brms1l |
| Gadd45a | Apbb3 | Spry1 | Meiob | Tnfsf13 | Atp6v0e2 |
| Ccn2 | Plekhg5 | Bhlhe40 | Gpr39 | Sccpdh | Foxa2 |
| Adamts1 | Fzd1 | Xkr8 | Xiap | Car1 | Plekhf1 |
| Cry2 | Poc5 | Pcyt1a | Svil | Gls | Pdcd5 |
| Ankrd33b | Nle1 | Rnf125 | Hnf4g | Pdgfrb | Oaf |
| Sdc4 | Trmt61a | Agt | Dtx4 | Ppp1r14a | Cbln3 |
| Itpr1 | LOC500948 | Ppp1r3c | Cryl1 | Dynlt1 | Dph2 |
| Fmo3 | Nop2 | Erf | Irf6 | Capg | Med22 |
| Nipal1 | Fndc3b | Ccnb2 | Phf11 | Fdps | LOC682870 |
| Limk2 | Gpd2 | Fus | Tmem159 | LOC691141 | Tstd2 |
| Asl | Flad1 | RGD1564804 | Fbxl15 | Slc38a9 | Htatip2 |
| Zbtb16 | Serpina5 | Srsf5 | Pop1 | Metap1d | Ears2 |
| Ppara | Akr1c14 | Eln | Mfhas1 | Atp23 | Abhd8 |
| Ppcs | Mpzl3 | Crebrf | Tprn | Csrp1 | Pex26 |
| Bach1 | Mrm1 | Pttg1 | Nrep | Twf2 | RGD1564171 |
| Slc16a10 | Niban1 | Pcsk6 | Tlr4 | Rhoh | Taok3 |
| Nr1d1 | Dusp16 | Cdkn3 | Depp1 | Pnrc1 | Ntf3 |
| Dact2 | Gas5 | Bcl3 | Bin2 | Pdk4 | Parp14 |
| Slc7a2 | LOC100134871 | Dnase2b | Steap3 | Marveld1 | Ubxn2b |
| Cpt1a | Aspg | Mrps31 | Dusp19 | Tra2a | Nat2 |
| Nnmt | Enpp2 | Gatd1 | Rps6 | Cml2 | Parp16 |

| | | | | | |
|---|---|---|---|---|---|
| Abhd15 | Ndufaf4 | Zfp622 | Bivm | Akr1d1 | Traf2 |
| Col4a4 | Med15 | Calcoco1 | P4ha1 | Mfsd13a | Giot1 |
| Fv1 | Baiap2 | Mrpl30 | Baat | Fdft1 | Ncald |
| Lpin2 | Slc22a25 | Cox11 | Lingo4 | LOC498276 | Hamp |
| Gpr146 | Exosc5 | Prkaa1 | Mblac2 | Slc46a1 | Oas1b |
| Klf16 | Faap20 | Chst13 | St3gal6 | Tcea3 | Rsrp1 |
| Klf10 | Xbp1 | Errfi1 | Cd8a | Prf1 | Nabp1 |
| Slc25a22 | Dancr | Cyp2e1 | Adamts7 | Ckb | Hsp90aa1 |
| Sbk1 | Per2 | Pex16 | Stx2 | Echdc1 | Mir293 |
| Derl3 | Spata5 | Sorbs3 | Slc16a1 | Paqr7 | Tubb4b |
| Lnc-hc | Polr2f | Spata2L | Pard6b | Ttc8 | Jmjd4 |
| Igfbp2 | Fitm2 | Pfkfb2 | Arhgap29 | Gldc | Tcta |
| Ctif | Marchf6 | Yod1 | Dynll1 | Nfil3 | Eva1a |
| Slc25a15 | Tle1 | Pex11a | | Clock | Tnfaip2 |
| Slc43a1 | Pde4dip | Dazap1 | | RT1-DMb | Ccl24 |
| Slc20a1 | Cited4 | Taf5l | | Plin2 | Rbm14 |
| Mtmr7 | Ppp1r10 | Klf6 | | Uba7 | Maz |
| LOC498154 | Fkbp14 | Id1 | | Mir186 | Gfer |
| Mat1a | Pald1 | Gsta3 | | Tspan14 | Als2 |
| Nfkbiz | Spout1 | Ephx2 | | Akr1c1 | Mitd1 |
| Mtss1 | Clasrp | Nedd9 | | Akip1 | Hhex |
| Oat | Zfp7 | Hand2 | | Hspb1 | P2ry1 |
| Ttc7b | Myo1b | RT1-DMb | | Gbp2 | Kpna4 |
| Gdnf | Znhit2 | Marchf7 | | Ift172 | Odf3b |
| Klf15 | Mrnip | Pf4 | | Pter | Creb1 |
| Slc7a10 | Slc37a4 | Zfp629 | | Sez6 | Uros |
| Srsf4 | Noc3l | Epha2 | | Lap3 | Ints6 |
| Acer2 | Minpp1 | Rcn3 | | Lyc2 | Prickle1 |
| Nr1d2 | Cpamd8 | Tspan18 | | Zfp36l1 | Ppil1 |
| Nfkbia | Acot12 | Jmjd7 | | Rnf125 | Creld2 |
| Cirbp | B3galt1 | Bmyc | | Pdzrn3 | Gata4 |
| Kif21a | Cyp2c12 | Rdh16 | | Tank | Trim21 |
| Tef | Ier3 | Arl5b | | Fasn | Hps6 |
| Wee1 | Pnpla7 | Gins4 | | Slc28a2 | Smc6 |
| Nr1i2 | Fam126b | Sephs2 | | Adipor2 | Zfyve21 |
| Fzd5 | Ogfod1 | Paxip1 | | Cdk6 | Tmtc4 |
| Slc2a5 | Eif4ebp2 | Nupl2 | | Slc45a3 | Bambi |
| Inmt | Fam98c | Ngdn | | Samsn1 | Mob3b |
| Polg2 | Wdr35 | | | Carhsp1 | Gimap4 |
| H2ac1 | Chn2 | | | MGC93861 | Ctns |
| Syde2 | Cep85l | | | LOC100910973 | Bmf |
| Abcb1a | Mir27b | | | Jade2 | Pacsin3 |
| Enc1 | Pus7 | | | Arrdc1 | Stk38l |
| Gpcpd1 | Timm8a1 | | | Loxl4 | Tmem51 |
| Dixdc1 | Tnk2 | | | Abhd5 | Fam219b |
| Nceh1 | Prrg1 | | | Thap4 | Zfp46 |
| Herpud1 | Irgq | | | Il18bp | Slc35a5 |
| Cndp1 | Slc38a4 | | | Sh3yl1 | Aktip |
| Il1rn | Kdsr | | | Mllt3 | Ror1 |
| Foxo1 | LOC497899 | | | Nipa1 | Rsl1d1l1 |
| Cyth1 | Hs3st3b1 | | | Fam89b | Zfp2 |
| Fem1a | Lin7c | | | Gpr182 | Zscan21 |
| Tgfb3 | Slc25a28 | | | Serpinb6b | Nde1 |
| Aass | Recql5 | | | Iffo2 | Map7d1 |
| Cd320 | Wdr77 | | | Ubd | Rassf3 |
| Chd7 | Kyat1 | | | Kxd1 | Meis1 |
| Tent4a | Mir5132 | | | Rasd1 | Zhx2 |
| Synj2 | Kif13a | | | Fgr | Eef2kmt |
| H1f4 | Slc25a26 | | | Mnda | Hmgxb4 |

| | |
|---|---|
| Rrs1 | Amdhd1 |
| Vcam1 | Lmbrd2 |
| Gpd1l | Mdm2 |
| Timp1 | Nob1 |
| B4galnt1 | Nifk |
| Nampt | Acot4 |
| Baiap2l1 | Elovl2 |
| Mgp | Acot3 |
| H6pd | Fhit |
| RGD1309748 | Pde4b |
| Bnip3 | RGD621098 |
| Pla2g15 | Fn3k |
| Mrps10 | Tmem220 |
| Abcb1b | Heatr1 |
| Abcg5 | Adck1 |
| Ifrd2 | LOC498368 |
| Epas1 | Prkce |
| Klf13 | Nudt22 |
| Mctp2 | Marveld2 |
| Arid5b | Cebpd |
| Gch1 | Cpz |
| Ces2c | Xrcc1 |
| Taf1d | Fam160a1 |
| Ptdss2 | LOC100365008 |
| Atp1b1 | Ankrd46 |
| Pak1ip1 | St3gal5 |
| Pxdc1 | Rapgef5 |
| Nrbp2 | LOC680200 |
| Arg1 | Gnl2 |
| Cyp3a23/3a1 | Fen1 |
| Mocs1 | Cyp3a9 |
| Zfp867 | Gpr137b |
| Nostrin | Dgka |
| Azin1 | Parp6 |
| Lrrfip1 | Utp15 |
| Wasl | Eml5 |
| Slc49a4 | Apex1 |
| | Ptn |
| | Slc4a1ap |
| | Abhd2 |
| | Smim22 |
| | Nop14 |
| | Fam174a |
| | Ltv1 |
| | Cyhr1 |
| | Tspyl2 |
| | Gpatch2 |
| | Rnf121 |
| | Trdmt1 |
| | Fam214b |
| | Bcl2l12 |
| | Endog |
| | Tex30 |
| | Bmp10 |
| | Fam76a |
| | Rab11fip2 |
| | Ctsl |
| | Tbc1d24 |
| | Ccdc51 |

| | |
|---|---|
| Crip1 | Ahsa2 |
| Fmo5 | Plxna2 |
| Rap2b | Irs1 |
| Stx18 | Zfyve19 |
| Wdr6 | Ska2 |
| Cryab | Snx16 |
| Slc17a3 | Cep44 |
| Otub2 | Zfp133 |
| Insig1 | Ormdl3 |
| Cks2 | Cyp2f4 |
| Stat1 | Rbm4b |
| Ociad2 | Stx17 |
| Tap1 | Plk2 |
| Sycp3 | Nfic |
| Rtn4 | Tbck |
| Oaz2 | Lto1 |
| Znrf2 | Cobl |
| Cmpk2 | Exosc3 |
| Fcgr3a | Rbm5 |
| Trim5 | Phf7 |
| Rb1 | Hist1h4m |
| Frmd8 | Tcf3 |
| Itga6 | Rbp7 |
| Erc2 | Eef1akmt1 |
| | Tmem186 |
| | Slc29a1 |
| | Jpt2 |
| | Unc119b |
| | Nt5c |
| | Gprc5c |
| | Nup43 |
| | Etfbkmt |
| | Hist1h2an |
| | Mrpl49 |
| | Gne |
| | Pms1 |
| | Alg2 |
| | Nuak2 |
| | Aifm2 |
| | Snx24 |
| | Lama3 |
| | Iqcb1 |
| | Sqor |
| | Mmp14 |
| | Dcun1d3 |
| | UST4r |
| | Foxk2 |
| | Zfp772 |
| | Rnf34 |
| | Lin7a |
| | Hmces |
| | Ttc9c |
| | Hspa14 |
| | Acap3 |
| | Kifbp |
| | Zfp444 |
| | Slc10a5 |
| | Mindy1 |
| | Plcb3 |

| |
|---|
| Zbtb26 |
| Cyp4a1 |
| Aqp11 |
| Dip2a |

| |
|---|
| Tnfsf10 |
| Gtf3c5 |
| Parp2 |
| Ints13 |
| Tmem50b |
| Tsen15 |
| Senp7 |
| Zfp455 |
| Dcaf7 |
| Ccndbp1 |
| Igfals |
| Tada3 |
| Mturn |
| Prxl2c |
| Zfp68 |
| Exo5 |
| Rwdd2b |
| Fem1b |
| Phtf1 |
| Pmm2 |
| Foxa1 |
| Dnajc28 |
| Fzd8 |
| Zfp35 |
| Zfp865 |
| Znfx1 |
| Hipk2 |
| Hist1h2bd |
| Ankrd54 |
| Zfp213 |
| Slc7a5 |
| Tcaim |
| Eno3 |
| Psph |
| Amz2 |
| Nuak1 |
| Fabp3 |
| Hspa8 |
| Ccdc117 |
| Abcg3 |
| Aagab |
| Stard5 |
| Mepce |
| Usp20 |
| Rnf43 |
| LOC100125367 |
| Rxrg |
| Ier2 |
| Phf11b |
| Rbm10 |
| Parp9 |
| Mrpl34 |
| Zfp52 |
| Gspt1 |
| Hdac5 |
| Reg4 |
| Upf2 |
| Rprd1a |
| Rita1 |

| |
|---|
| Ndufa1 |
| Msrb2 |
| Deaf1 |
| Trmt11 |
| Fas |
| Arl14ep |
| Ttc5 |
| Osbpl7 |
| Ctu1 |
| Hexim1 |
| Pdp1 |
| Dennd11 |
| Magix |
| Zfp322a |
| Rfc1 |
| Slc25a30 |
| Cyp4f6 |
| Arhgef40 |
| Cxxc1 |
| Plekha8 |
| Trip10 |
| Atp5pb |
| Rab11fip4 |
| Rnpc3 |
| Ing4 |
| Usf1 |
| Nup35 |
| Mphosph8 |
| Rbmxrtl |
| B4galt3 |
| Kbtbd7 |
| Ppil3 |
| Akr1c2 |
| Crls1 |
| Ifi44 |
| Thumpd3-as1 |
| Taf12 |
| Sfpq |
| Ddx17 |
| Itsn1 |
| RGD1566325 |
| Ccl27 |
| Fam214a |
| Kbtbd3 |
| Nod1 |
| Acnat2 |
| Ocln |
| Atf5 |
| Them6 |
| Thumpd1 |
| Mocs2 |
| Trim39 |
| Klhl42 |
| Mir3064 |
| Mif4gd |
| Slc10a3 |
| Fam118b |
| Hikeshi |
| Arhgap24 |

| |
|---|
| Zfp51 |
| Prkci |
| Dlg3 |
| Clic4 |
| Zfp467 |
